# Evading resistance to gene drives

**DOI:** 10.1101/2020.08.27.270611

**Authors:** Richard Gomulkiewicz, Micki L. Thies, James J. Bull

## Abstract

Gene drives offer the possibility of altering and even suppressing wild populations of countless plant and animal species, and CRISPR technology now provides the technical feasibility of engineering them. However, population-suppression gene drives are prone to select resistance, should it arise. Here we develop mathematical and computational models to identify conditions under which suppression drives will evade resistance, even if resistance is present initially. Previous models assumed resistance is allelic to the drive. We relax this assumption and show that linkage between the resistance and drive loci is critical to the evolution of resistance and that evolution of resistance requires (negative) linkage disequilibrium between the two loci. When the two loci are unlinked or only partially so, a suppression drive that causes limited inviability can evolve to fixation while causing only a minor increase in resistance frequency. Once fixed, the drive allele no longer selects resistance. Our analyses suggest that among gene drives that cause moderate suppression, toxin-antidote systems are less apt to select for resistance than homing drives. Single drives of moderate effect might cause only moderate population suppression, but multiple drives (perhaps delivered sequentially) would allow arbitrary levels of suppression. The most favorable case for evolution of resistance appears to be with suppression homing drives in which resistance is dominant and fully suppresses transmission distortion; partial suppression by resistance heterozygotes or recessive resistance are less prone to resistance evolution. Given that it is now possible to engineer CRISPR-based gene drives capable of circumventing allelic resistance, this design may allow for the engineering of suppression gene drives that are effectively resistance-proof.

## Introduction

The ability to engineer gene drives has recently become feasible for countless species, especially insects, many of which carry disease or eat crops (Gantz and Bier 2015; Champer *et al.* 2016; Kyrou *et al.* 2018; Oberhofer *et al.* 2019). Gene drives are genetic elements that use biased segregation or directed killing of alternative alleles to give themselves an ‘unfair’ evolutionary advantage, possibly to the detriment of their carriers. One possible use of engineered drives is to suppress populations (’suppression’ drives), to reduce or even eradicate an entire species (Hamilton 1967; Lyttle 1977; Burt 2003). Perhaps the major remaining biological hurdle in successfully implementing suppression drives is avoiding the evolution of resistance. In the early history of gene drive research, of both natural and experimental examples, resistance was the outstanding cause of the persistence of gene drives that would otherwise have extinguished populations or suppressed them to far greater levels (Lewontin and Dunn 1960; *Sandler et al.* 1959; Sandler and Hiraizumi 1959; Lyttle 1979, 1981). Of course, examples from nature are necessarily devoid of past extinctions, so only experimental work can truly reflect on the *a priori* likelihood of resistance evolving to suppress a drive that will cause extinction.

Gene drive engineering has been profoundly facilitated by CRISPR technology (Gantz and Bier 2015; Champer *et al.* 2016; Oberhofer *et al.* 2019). CRISPR uses an RNA-directed nuclease to cut specific sites in DNA, enabling a variety of gene drive designs from homing endonucleases that operate by segregation distortion to killer-rescue and toxin-antidote systems that destroy sensitive alleles. The most obvious form of resistance to CRISPR-based drives is mutation in the target sequence at which the nuclease cuts (Burt 2003; Unckless *et al.* 2015, 2017; Champer *et al.* 2017). However, contrary to early suspicions that all possible target sites would be prone to such mutations (e.g., Drury *et al.* 2017), at least one essential genomic site has been identified in mosquitoes that is intolerant of mutations and thus appears to escape this problem (Kyrou *et al.* 2018). Furthermore, CRISPR-based gene drives can be designed to cut at multiple sites in the same gene (any of which suffice to allow the drive to function), thus ameliorating most concerns about the inevitability of target-site resistance (Oberhofer *et al.* 2018; *Champer et al.* 2018, 2020c). Other forms of resistance to CRISPR-based gene drives are theoretically possible, but it is too early to tell whether they will prove to interfere with implementations: inbreeding, proteins that block CRISPR complex activity, and suppressors of CRISPR gene expression (Stanley and Maxwell 2018; *Bull et al.* 2019). Experimental work by Lyttle (1979, 1981) using a natural gene drive mechanism raise the spectre of diverse and unpredictable forms of resistance to strongly suppressing gene drives, so resistance evolution is intrinsically plausible from our limited evidence.

All forms of resistance are not necessarily equal. Target-site resistance differs from other potential types of resistance in one important respect: linkage to the drive. For gene drives that operate as homing endonucleases, target-site resistance is allelic, segregating opposite the drive itself (Unckless *et al.* 2015, 2017; Drury *et al.* 2017). Many of the possible alternative types of resistance will not be allelic to the drive, they may even be completely unlinked (Lyttle 1981; Champer *et al.* 2019). The purpose here is to understand the evolution of resistance to a population-suppressing gene drive when the resistance is potentially unlinked to the drive. Our results show that loose linkage between the resistance and drive loci works against the evolution of resistance for a wide range of conditions. This realization allows us to identify strategies that may avoid the evolution of resistance to gene drives and still – by the sequential introduction of multiple drives – allow eventual population suppression to arbitrary levels.

## Models and Results

We formulate deterministic population genetic models with two and three loci to study the joint evolution of gene drive and resistance. Exact analytical results are derived for relatively simple cases to understand the processes affecting the evolution of resistance. The derivations, detailed in the Appendix, track haplotype frequencies after transmission and selection, with expressions then mathematically transformed into allele frequencies and linkage disequilibria. Computation employing mostly C programs (see Supplemental File S1), some with R (R Core Team 2019) was used to study more complex cases.

### Population genetics with non-allelic resistance

A two-locus model with selection in haploids (gametes) illustrates some key properties of selection on and evolution of gene drives. The setup for this model is extreme for purposes of illustration: segregation distortion is assumed to be 100%, the two loci are fully unlinked, and resistance is dominant, complete, and has no fitness cost; analyses in the Appendix are more permissive (e.g., imperfect distortion, incomplete dominance, linkage). Segregation distortion operates at the *A*/*a* locus, resistance to segregation distortion operates at the *R*/*r* locus (Table 1).

**Table 1.**
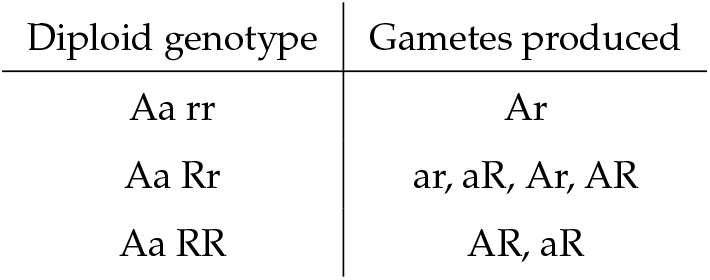
Segregation ratios for the three possible genotypes that are heterozygous at the drive locus. Only the genotype of the top row experiences segregation distortion, as any genome with allele *R* suppresses distortion. All of the gamete types listed for a diploid are produced in equal abundance; the double heterozygote produces all 4 in equal abundance because of the absence of linkage between the two loci. Segregation is Mendelian for the 6 other possible diploid genotypes not shown here – those involving *AA* and *aa* homozygotes at the drive locus.

Fitness effects are expressed in the gamete stage, with only the gene drive allele having an effect (Table 2). (The Appendix includes analysis of a case of non-gametic selection, namely, a recessive lethal gene drive.) The following sections explain in stepwise fashion how unlinked resistance is selected.

**Table 2.** Haplotype (gamete) fitnesses. *s*_*A*_ is the coefficient of selection agains the drive allele. In this model, fitnesses are assigned to gametes rather than diploids.

| Gamete type | Fitness |
| --- | --- |
| ar | 1 |
| Ar | $1 - s_A$ |
| aR | 1 |
| AR | $1 - s_A$ |

### The non-drive allele is depressed by segregation distortion but bolstered by its fitness advantage

In the absence of resistance alleles, the net effect of each life cycle is to depress the frequency of the non-drive allele *a*. But this net effect is the outcome of the opposing effects of segregation distortion and gametic fitness. Assuming transmission occurs prior to selection against the drive allele *A*, the frequency of allele *a* after transmission but before selection 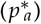 as a function of its frequency at the start of the generation (*p*_*a*_) is

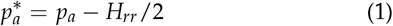

where *H*_*rr*_ is the frequency of diploid genotypes that are resistance-free and heterozygous for the drive (*Aarr*). The term *H*_*rr*_/2 describes the depression of allele *a* from 100% segregation distortion. Note that resistance will do nothing to blunt this negative effect, other than perhaps to reduce *H*_*rr*_.

Selection against the drive subsequently counters this. The frequency of *a* is increased from 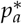 by the factor 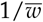,

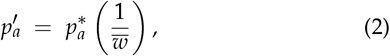

where 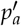 denotes the frequency of *a* at the start of the next generation and 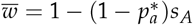 is mean fitness, necessarily less than 1 when the drive allele is present. The term in parentheses in (2) describes the gain of allele *a* from its fitness advantage over *A*. The loss in (1) is always larger than the gain (2) when the drive allele is spreading. However, it is the separation of these effects that is critical to understanding the evolution of non-allelic resistance.

### Resistance has no advantage in the transmission stage

This point is perhaps the most surprising, but is easy to understand. Considering the resistance allele *R* before and after the segregation distortion stage, there is no change in its frequency:

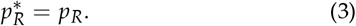

This result may be understood intuitively from the fact that every genotype with *R* segregates normally. Even in the case of the *AaRr* double heterozygote, the lack of segregation distortion (recall *R* is assumed dominant) means that *R* will on average occur in half the gametes (regardless of the haplotypes that formed the double heterozygote; see Table 1), the same frequency as in the parent. This holds even if the loci are linked. Segregation distortion occurs only in the *Aarr* dipoid, and all gametes carry *r*, so again there is no change in its frequency.

Given that neither *R* nor *r* has an effect on gametic fitness, there is no intrinsic advantage that one allele has over the other during the transmission stage. However, the relative benefit of resistance changes when we consider associations among loci, as next.

### Synergy between resistance and fitness of the non-drive allele

The foregoing two sections establish that the non-drive allele *a* gains in survival over *A* each generation but loses potentially more from segregation distortion. By itself, the resistance allele has no fitness advantage and also does not benefit by suppressing distortion at the other locus. Both outcomes change when we consider resistance combined with the fitness advantage of the non-drive allele. The two alleles synergize when they occur in the same genotype. When associated with *R*, the *a* allele is emancipated from its loss in segregation distortion because segregation distortion is abolished; now its only deviation from neutrality is the gain due to its intrinsic fitness advantage in gametes. In turn, allele *R* gains by segregating with *a*, experiencing the fitness advantage *a* has in gametes. Thus, both *R* and *a* benefit from their co-occurrence, each allele providing a benefit to the other.

For this association to provide a net benefit across the population, the alleles of the different loci must be statistically associated. In population genetics terms, there must be (negative) linkage disequilibrium, which can be defined as

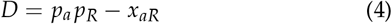

where *x*_*aR*_ is the frequency of gametes that contain both *a* and *R*; *D* < 0 indicates that *a* and *R* occur together more frequently than at random. Note that *D* is bounded below by the larger of −(1 − *p*_*a*_)*p*_*R*_ and −*p*_*a*_ (1 − *p*_*R*_) (e.g., Crow and Kimura 1970). This lower bound to the strength of the association between *a* and *R* approaches zero in particular as the frequency of the drive allele *A* approaches 1.

During transmission, free recombination halves the magnitude of *D* but, separately, segregation distortion forces the association between *a* and *R* closer. Specifically, the disequilibrium after transmission, *D*\*, becomes

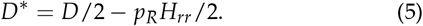

The subtracted term shows that the amount of association created by gene drive increases with the frequencies of resistance-free drive heterozygotes (*H*_*rr*_) and the resistance allele itself. If *D* is initially positive, recombination and segregation distortion work in concert to reduce it. Once *D* is negative, the two forces work in opposition with segregation distortion exaggerating and recombination eroding the association between *a* and *R*.

The statistical association between *a* and *R* has consequences for *R* when *a* is selected. After selection its frequency becomes

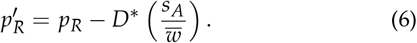

This result shows that negative linkage disequilibrium (*D*\* < 0) is required for *R* to rise in frequency – that *a* and *R* must exist together in the same gamete more often than random. Thus, non-allelic resistance increases by genetic hitchhiking (Maynard Smith and Haigh 1974).

To summarize, this simple model highlights three main features: Segregation distortion has no direct effect on resistance (unless allelic to the drive), segregation distortion directly promotes a tighter statistical association between *a* and *R*, and alleles *a* and *R* must co-occur in gametes more often than at random (i.e., negative linkage disequilibrium is required) for resistance to rise in frequency via hitchhiking.

Below and in the Appendix, we consider more complex scenarios with incomplete segregation distortion, partial dominance, linkage between drive and resistance loci, costly resistance, sex-limited and toxin-antidote gene drives, and diploid selection, including lethal recessive drives. These models show that the scope for resistance evolution increases with more effective segregation distortion, closer linkage, and less costly resistance. Despite their quantitative influences on the evolution of resistance, those features do not affect the three key features emphasized above.

### Evading resistance evolution is possible and practical

The preceding results facilitate developing protocols for suppression drives that diminish the evolution of resistance. Prior to exploring whether such protocols are theoretically feasible, we offer several points to serve as intuitive guiding principles.

1. Allelic resistance should be avoided. Resistance is least prone to evolve when weakly linked to the drive. Some designs using CRISPR homing endonucleases can greatly limit the opportunity for allelic resistance.
2. Resistance may impose a cost, independent of the drive itself. Any cost will work against resistance evolution.
3. The drive should be designed to evolve to fixation. Even if resistance arises, it cannot maintain the required linkage disequilibrium once the drive allele fixes, so it will cease to function as the drive sweeps to high levels. Persistent heterozygosity of a drive allele provides an indefinite opportunity for resistance to reverse the drive at any time prior to species extinction. Drive fixation precludes future reversals.
4. The drive homozygote should be viable and fertile, with a substantial but not extreme reduction (~10%–20%; see Fig. 1 below) in fitness. A modest fitness effect not only facilitates drive fixation (point 3, above) but also avoids the spread of resistance alleles with a high fitness cost.

**Figure 1.**
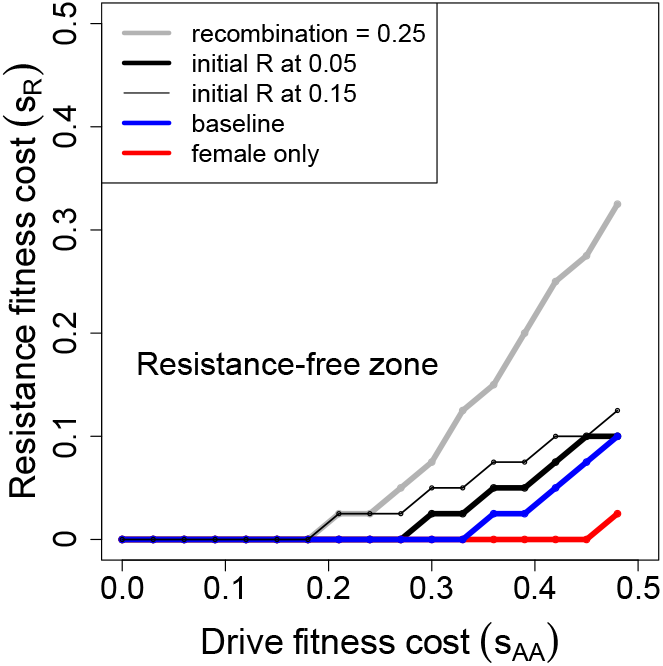
Evolution of (dominant) resistance to a male-only drive: numerical results. The drive allele can evolve to fixation despite the presence of resistance alleles in the population, depending on the magnitude of the suppression (fitness of drive homozygotes) and on the cost of resistance. Gene drives evolve to fixation above and to the left of the curves, in the ‘resistance-free’ zone, at which point they are no longer subject to suppression. The baseline trial (blue) assumes drive is 100% efficient, the drive allele starts at frequency 0.005, resistance at 0.015. The drive allele impairs fitness of homozygotes of both sexes, their fitness being 1 − *s*_*AA*_. Resistance to the drive’s segregation distortion is unlinked (except for the dashed gray curve), dominant, complete, and impairs fitness of *Rr* and *RR* genotypes as 1 − *s*_*R*_. For the baseline case, even cost-free resistance does not evolve until *s*_*AA*_ > 0.3, and at higher values of *s*_*AA*_, resistance evolves and blocks fixation of the drive allele only if cost of resistance (*s*_*R*_) is sufficiently low. Other curves deviate from the baseline case in single respects: (red) the fitness effect of drive homozygotes *s*_*AA*_ is experienced only in females; (thick black) the initial frequency of resistance is increased to 0.05; (thin black) the initial frequency of resistance is increased to 0.15; (gray) the drive and resistance loci are linked with a recombination rate of 0.25. Although not shown, the line is flat out to *s*_*AA*_ = 0.54 if the fitness effect *s*_*AA*_ is assigned only to males instead of to females. All trials were run at least 700 generations; values of *s*_*AA*_ were incremented by 0.03, *s*_*R*_ by 0.025. Trials initiated *R* and *A* in separate individuals (negative disequilibrium), but limited simulations suggested the curves are not sensitive to this aspect of initial conditions.

### Imperfect distortion, partially dominant resistance, and genetic linkage

The analytical results above assume segregation distortion is 100%, resistance is dominant and complete, and that the drive and resistance loci are unlinked. Relaxing these assumptions has the following consequences. (See Appendix for all derivations.)

First, let the segregation distortion for resistance genotype *G* be denoted by *δ*_*G*_, where *G* = *RR*, *Rr*, and 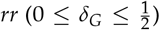. In the Appendix, we show that, following transmission, the non-drive allele frequency is

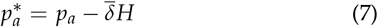

where *H* is the total frequency of drive heterozygotes *Aa* and 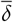 is the average distortion across all three resistance locus genotypes (see eq. 11). Equation (7) reduces to (1) if resistance is dominant and complete (*δ*_*Rr*_ = *δ*_*RR*_ = 0) and if distortion in the absence of resistance is 100% 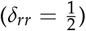.

Second, transmission can directly change the frequency of the resistance allele if heterozygous resistance is incomplete (*δ*_*Rr*_ > 0). Assuming random mating, this frequency is

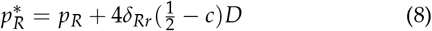

where *c* is the recombination rate between the drive and resistance loci and *δ*_*Rr*_ is the drive segregation distortion allowed by resistance heterozygotes (see Appendix, eq. 14). Equation (8) shows that transmission will have no effect on the frequency of resistance (i.e., 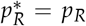; cf. eq. 3) unless three conditions are simultaneously fulfilled: the loci must be (i) linked 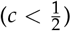, (ii) in linkage disequilibrium (*D* ≠ 0), and (iii) resistance heterozygotes allow some segregation distortion (*δ*_*Rr*_ > 0). Note too from (8) that 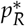 is not influenced by segregation distortion in *rr* or *RR* resistance genotypes; this is a completely general result (see the Appendix). Since *D* can be positive or negative, (8) shows that transmission might increase or decrease the frequency of *R*. As above, transmission can change *D* itself. Assuming random mating, the disequilibrium after transmission becomes

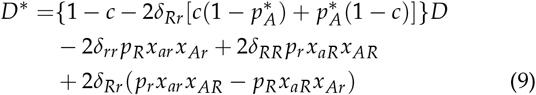

where 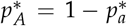 is the post-transmission frequency of the drive allele *A*, *x*_*ij*_ is the pre-transmission frequency of haplotypes with drive allele *i* and resistance allele *j*, and *δ*_*rr*_ and *δ*_*RR*_ are the respective segregation distortions of wild type and resistance homozygotes (see Appendix, eq. 16). Note that (9) reduces to (5) when drive and resistance are unlinked 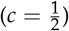, resistance is complete and dominant (*δ*_*RR*_ = *δ*_*Rr*_ = 0) and segregation distortion is maximal in the absence of resistance 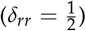.

Equation (9) shows that partial resistance can affect disequilibrium’s rate of decay (i.e., the factor in braces that multiplies *D*) as well as the generation of new disequilibrium (bottom two lines of the expression). Only incomplete resistance in *Rr* heterezygotes can influence the decay rate. If heterozygous resistance is complete *δ*_*Rr*_ = 0, disequilibrium decays at the usual rate under random mating, namely, by the factor 1 − *c*. However, if the resistance is partial (*δ*_*Rr*_ > 0) then disequilibrium decays more rapidly since the term in square brackets is always positive. If *D* = 0, the second two lines of (9) show that segregation distortion by all resistance genotypes can contribute positively or negatively to the post-transmission disequilibrium. Our numerical results (see below) suggest that the negative influences dominate over time, reducing and eventually leading to generations of negative *D* values. As a consequence (see eq. 8), transmission with imperfect heterozygous resistance will tend to reduce the frequency of resistance directly. Since transmission does not affect the frequency of perfect and dominant resistance (eq. 3), this suggests that gene drives engineered to prevent dominant resistance from evolving would also prevent the spread of partially dominant and recessive resistance.

### Numerical studies

We offer numerical analyses of models that fulfil many of the preceding points (code available in Supplemental File S1). These numerical models assign fitness effects to diploids, appropriate because diploid fitness effects would accrue to current implementations. (i) The drive is expressed in one sex (males) or both sexes. (ii) The drive operates with 100% segregation distortion. (iii) The drive allele depresses the fitness of drive homozygotes only (to 1 − *s*_*AA*_). A value of *s*_*AA*_ near 1 reflects a drive’s strong population suppression effect, mild to moderate otherwise. In some implementations, the fitness effect accrues to drive homozygotes of both sexes; in others just to females. (iv) The resistance allele (*R*) fully suppresses segregation distortion of the drive – the most favorable case for its evolution. *R* is assumed dominant in some models, recessive in others. The fitness of *R* carriers is also varied. Most models analyzed are of homing drives, but one form of toxin-antidote system is modeled at the end.

### Male homing drive, dominant resistance

A suppression drive may be engineered to operate/drive in one sex even if the fitness effect on homozygotes is realized in both sexes. A one-sex drive cannot evolve to fixation (Prout 1953; Bruck 1957), the strength of population suppression being potentially milder than with a 2-sex drive. It might thus seem that one-sex drive is intrinsically less prone to favor resistance. Figure 1 shows long term (quasi-equilibrium) results of trials in which the drive homozygote fitness (1 − *s*_*AA*_) is varied along with the cost of resistance, *s*_*R*_; resistance is dominant, and the fitness associated with resistance (1 − *s*_*R*_) applies to heterozygotes (*Rr*) as well as homozygotes (*RR*). Most trials illustrated assume resistance is unlinked to the drive.

In the absence of resistance, the drive evolves to fixation (loss of *a*) provided *s*_*AA*_ ≤ 0.5. It is immediately apparent that drives with even substantial homozygous fitness effects (up to *s*_*AA*_ = 0.35) evolve to fixation, even when resistance imposes no fitness cost. But it is also seen that the outcome of evolution depends on the starting frequency of resistance: as the initial frequency of *R* is increased from 0.015 to 0.05 to 0.15 (blue, thick black, thin black), the drive becomes more susceptible to evolution of resistance – meaning that resistance prevents fixation of the drive. Arranging the drive’s fitness impairment to just females offers a considerable extension of the zone in which resistance fails to evolve (red curve). As foreshadowed from the analytical models, tightening the linkage of the two loci to 0.25 substantially facilitates the evolution of resistance, but there remains a considerable range of *s*_*AA*_ for which resistance does not evolve.

Frequency dynamics are shown in Figure 2 for a single trial. Resistance experiences a modest gain during gene drive evolution (red curve), reflecting the temporary evolution of negative linkage disequilibrium (both gray curves, separate for males and females). The increase is not enough to halt drive fixation but would be permanent in the absence of a cost to resistance. Costless resistance would therefore ‘ratchet’ itself up during the process, even when it does not ultimately block fixation of the drive, but any cost would continually bring resistance down between gene drive introductions.

**Figure 2.**
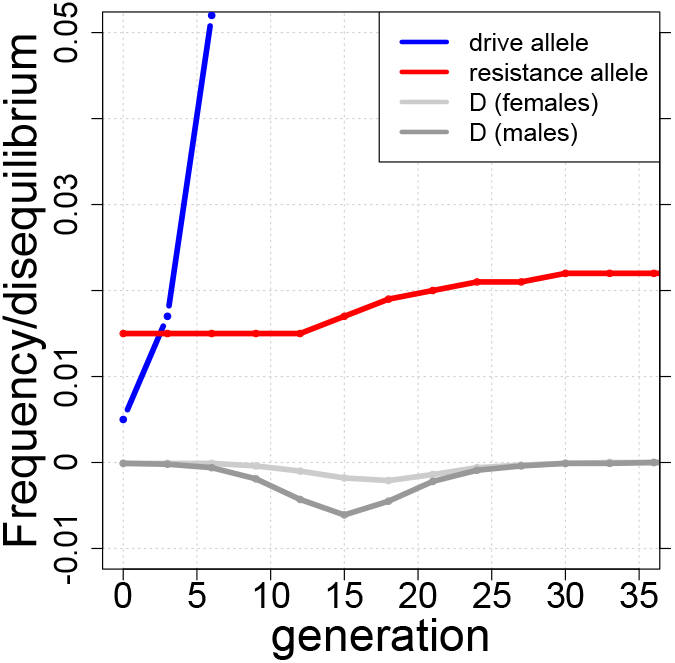
Dynamics of gene drive (blue), resistance (red), and linkage disequilibrium (gray) in a single trial. Linkage disequilibrium (D) is more pronounced in males (the sex experiencing the drive). The resistance allele (red) rises via indirect response to selection against the drive allele *A* during the transient submersion of disequilibrium below 0. For this trial, *s*_*AA*_ = 0.2, the resistance allele started at frequency 0.015 and entailed no cost (*s*_*R*_ = 0). This trial otherwise corresponds to the baseline case in Fig. 1.

The parameter space allowing resistance-free evolution of a homing male-drive is narrowed somewhat if fitness of the *Rr* heterozygote is intermediate between that of the two homozygotes (Fig. 3). The zone in which cost-free resistance does not evolve is necessarily not affected; the difference between the two cases is most pronounced with partial recombination (gray curve). In the region where cost-free resistance does evolve, resistance evolves more readily with semi-dominant fitness effects (here) than with dominant ones (Fig. 1).

**Figure 3.**
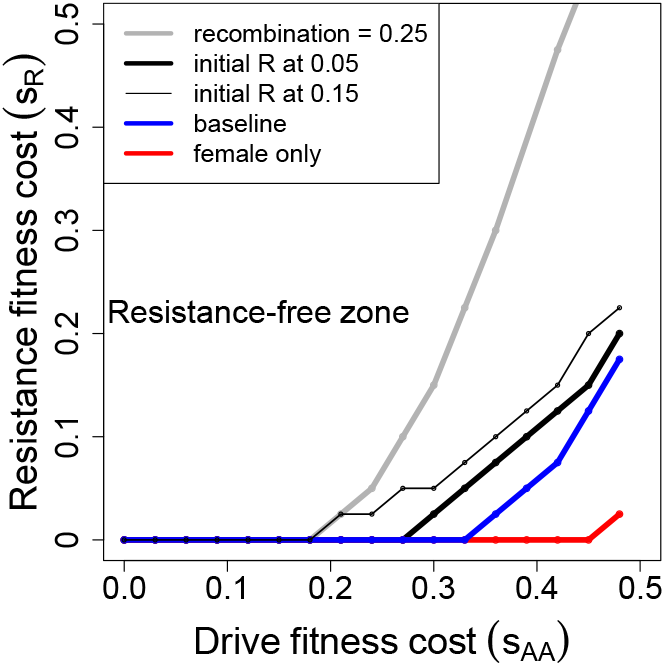
The evolution of dominant resistance is less hindered when the fitness effect of the resistance allele is additive (*Rr* has fitness 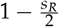, *RR* has fitness 1 − *s*_*R*_). Otherwise as in Fig. 1: the drive operates in males only.

### 2-sex homing drive, dominant resistance

2-sex drives are now easily implemented, so it is valuable to analyze that case for comparison to male-only drive (Fig. 4). The qualitative patterns are surprisingly similar up to the drive’s fitness effect of *s*_*AA*_ = 0.5 (compare Figs. 4 and 1; note that the axes’ scales differ). The main difference is that a 2-sex suppression drive allele can fix at higher *s*_*AA*_ values than can the male-only drive allele (1.0 versus 0.5), so there is more latitude for fixation of a 2-sex drive. However, the higher *s*_*AA*_ values more strongly select resistance, so a 2-sex drive does not obviously provide a more practical solution to resistance-free evolution than does a 1-sex drive.

**Figure 4.**
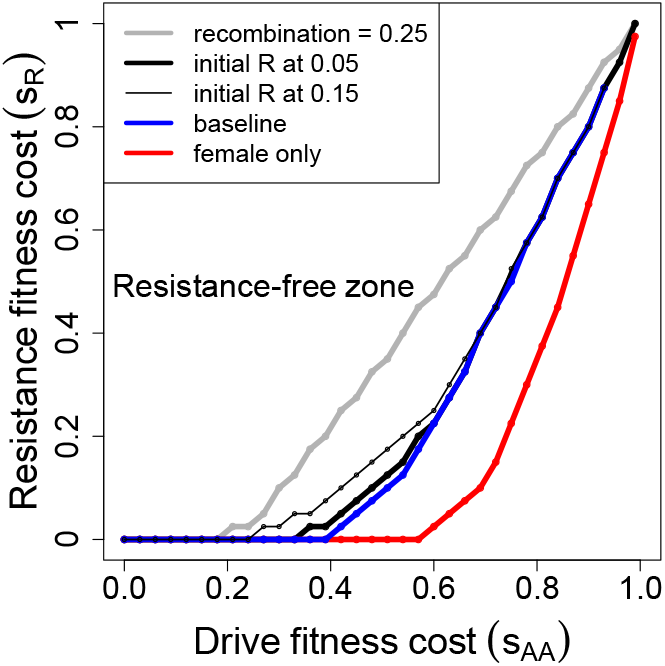
Evolution of dominant resistance to suppression drives that operate in both sexes. The overall patterns are qualitatively similar to those for male-only drives, except that 2-sex drives can fix for much higher values of *s*_*AA*_. Assumptions are otherwise the same as for Fig. 1.

### Male homing drive, recessive and partially dominant resistance

Negative linkage disequilibrium is essential to the selection of resistance, and the negative linkage disequilibrium is generated by resistance benefiting the drive-susceptible allele *a*. A reasonable conjecture is that dominance of the resistance allele will have a strong effect on evolution of resistance. To consider this possibility, the male-drive case of Figure 1 was studied for the case of recessive resistance – only the *RR* genotype suppressed drive, but the fitness effect of 1 − *s*_*R*_ applied to heterozygoes and homozygotes (Fig. 5). There is a profound effect of dominance versus recessivity of *R*, resistance being far less prone to evolve when it is recessive. Likewise, we found that partially dominant resistance was much less less likely to evolve than fully dominant resistance (results not shown). The engineer is likely not able to control the degree of dominance of resistance, but these results at least suggest that complete dominance is a worst-case scenario in anticipating the evolution of resistance.

**Figure 5.**
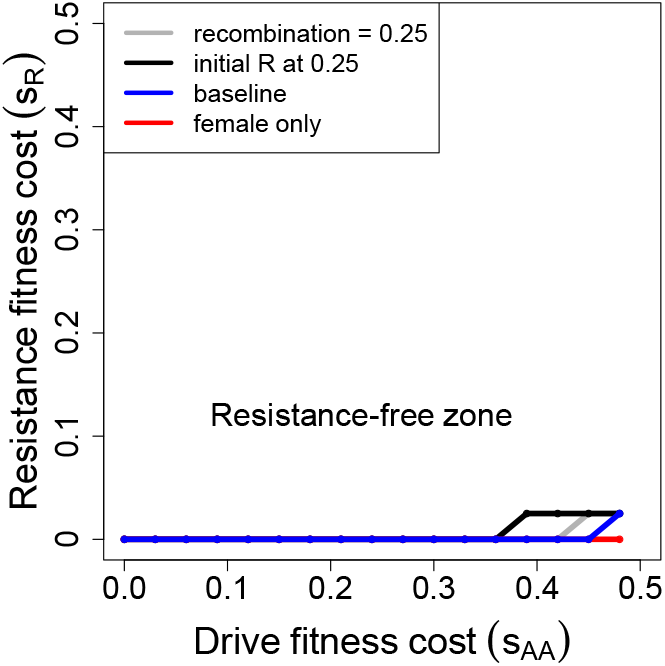
Evolution of recessive resistance to a male-only drive. Evolution of resistance is profoundly suppressed compared to the dominant-resistance case. Otherwise as in Fig. 1, except that only one (extreme) boost in initial frequency for *R* is considered, and even it has only a modest effect.

### Toxin-antidote drive, dominant resistance

Homing drives are highly efficient and have raised concerns about spread to unwanted target populations and species (Esvelt *et al.* 2014; Report 2016; Vella *et al.* 2017; Dhole *et al.* 2018). Some other types of drives pose less of a ‘spillover’ risk because they do not spread so easily, at least when rare. One class of ‘mild’ drives is comprised by killer-rescue or toxin-antidote systems (*Gould et al.* 2008; Marshall and Hay 2011; Oberhofer *et al.* 2019). The drive element in these systems has two functions. One function kills sensitive individuals; this function is distributed beyond individuals carrying the drive. The second function is to rescue carriers of the gene drive from the lethality.

A simple version of a toxin-antidote system is embodied by a recently invented design known as ‘Cleave-Rescue’ (*ClvR*) (*Oberhofer et al.* 2019). Using CRISPR technology, the *ClvR* element is engineered as a two-component system: (i) a Cas-9 nuclease targeted to destroy all native copies of an essential gene in the genome, and (ii) a recoded version of the essential gene that is protected against cleavage. The *ClvR* element – containing both components – may be inserted anywhere in the genome; it is easily engineered to be located at a site remote from and unlinked to the target gene, but ‘same-site’ designs are also possible (e.g., Champer *et al.* 2020a).

The *ClvR* element (denoted here as *C*, its absence being denoted *c*) spreads in an analogy to bootstrapping – it creates and disseminates a genetic ‘poison’ for which it provides the only antidote. This poison is merely the destruction of the target gene (the essential wild-type allele *T* is converted to a null allele *t*). As the null *t* alleles are introduced and spread in the population, they eventually form *tt* homozygotes which die unless the genome also carries at least one *C* allele (the antidote). Different variations of this theme can be engineered, but we will consider the case in which all genotypes have normal fitness except *cctt*, which dies.

While *C* is rare, *tt* is also very rare (under random mating), so the spread of *C* is slow (Oberhofer *et al.* 2020). If *C* imposes a cost to its carrier, this cost may be enough to prevent its ascendance unless introduced above a threshold frequency (*Oberhofer et al.* 2019). But as *C* becomes common, so does *tt*, and the benefit to *C* grows. Once *T* is extinguished, the killer property ceases to operate, and the population then evolves according to the fitnesses of the 3 possible genotypes at the *C* locus: *C* will not fix if *Cc* has higher fitness than *CC*, for example (all *cc* die).

For perspective, if there is no intrinsic fitness effect of *Cc* or *CC*, *allelic* resistance to destruction of *T* can evolve, but it does not necessarily cause the loss of *C*. In the extreme case of an allele *T*\* that is resistant to cleavage but suffers no fitness cost (all genotypes with *T*\* experience normal fitness), *T*\* will evolve at the same rate as *C*, the final frequencies seemingly in proportion to their initial frequencies. The endpoint of this process is loss of the wild-type (sensitive) *T* with ultimate fixation of *C* and/or *T*\* (simulation results not shown). The selective equivalence of *C* and *T*\* in this extreme case of no fitness effects is intuitive. A fitness difference between *C* and *T*\* would destroy that equivalence so that the one with higher fitness would prevail.

What about non-allelic resistance? We are especially interested in a parallel to the preceding analyses of homing drives that impose a perhaps modest fitness penalty on their carriers. Like homing drives, *ClvR* can be used for population suppression if it is inserted into and disrupts an important genomic region. This effect is different than the ‘toxic’ function which destroys the target gene. By inserting *ClvR* into the middle of an important gene, it now imposes a fitness cost on *Cc* and/or *CC*. *C* can potentially spread despite this cost, although it may need to be introduced above a threshold frequency (*Oberhofer et al.* 2019).

The evolution of resistance is no longer intuitive when *C* affects the fitness of its carriers and when resistance is non-allelic. First note that resistance to a *ClvR* system is merely a block to the ‘toxic’ property of *ClvR* – a block to the conversion of *T* to *t*. If *C* fully evades resistance evolution, then the population evolves to the complete loss of *T*. Once *t* is fixed, there is no further cleave activity on which selection for resistance may act. At this endpoint, *cc* becomes a universally lethal genotype, and if *Cc* has higher fitness than *CC*, the *C* locus will evolve to a stable polymorphism.

In the spirit of previous sections, we let *CC* genotypes have fitness 1 − *s*_*CC*_, and *Cc* genotypes have fitness 1, regardless of genotype at the target locus. *cc* genotypes have fitness 1 except in *tt* genotypes, when they are inviable. Resistance allele *R* is dominant in its blocking effect and in its fitness effect (1 − *s*_*R*_), paralleling the cases in Figs. 1 and 4. Fig. 6 shows the parameter space of *s*_*R*_ and *s*_*CC*_ that evades resistance evolution (for which *t* goes to fixation). For fitness effects above 0.2, the *ClvR* system is far less prone to evolve resistance than are the homing drive systems.

**Figure 6.**
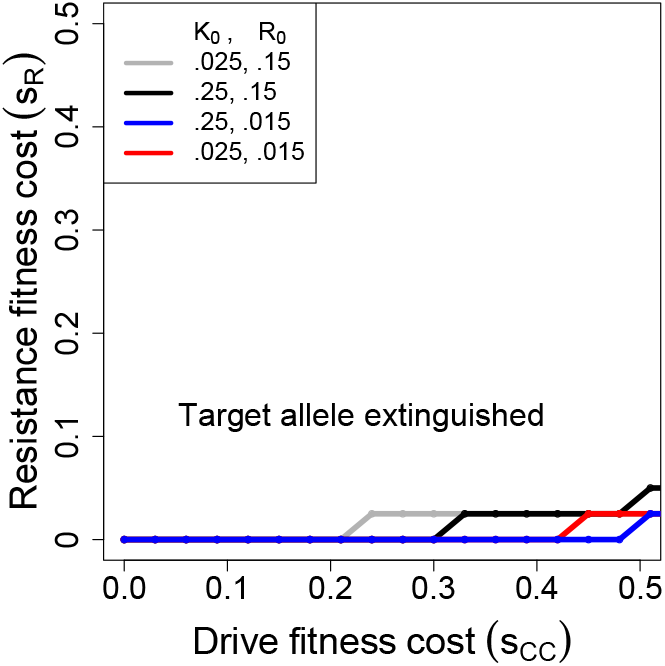
Evolution of dominant resistance to a toxin-antidote drive (*ClvR*) with no sex differences: resistance evolves only with a large fitness effect of the drive and low cost to resistance. Initial frequencies of the killer (K_0_) and resistance alleles (R_0_) are given in the key. The curves show the boundary of parameter values for which the wild-type target allele is fully converted by (*ClvR*) after 1,000 generations. *s*_*CC*_ refers to the fitness suppression of drive homozygotes (genotype *CC*), *s*_*R*_ to the fitness suppression of genotypes with one or two copies of the resistance allele.

## Discussion

In theory, gene drives offer an ability to quickly alter the genetics of populations, with two important uses: genetic modification or suppression of population numbers (Sandler and Novitski 1957; Hamilton 1967; Lyttle 1977; Burt 2003; Gould *et al.* 2006, 2008; Gould 2008). Modification drives facilitate the spread of genes without greatly affecting the population in other ways, although a modification drive might inadvertently impose a mild fitness cost on its carriers. Suppression drives can be used to permanently depress population densities, even to the point of extinction. Suppression can be simply a matter of creating a gene drive that has deleterious effects in drive homozygotes (Burt 2003), although other mechanisms are possible, such as sex ratio distortion (Hamilton 1967). CRISPR endonucleases now trivially enable the engineering of a type of drive known as a homing endonucleases (HEGs) that can be used for suppression or modification. For suppression, the drive is merely engineered to disrupt a gene of arbitrary importance to fitness, such that drive homozygotes are greatly impaired (Burt 2003).

Homing endonuclease genes work on the principle that the enzyme cuts a specific and unique DNA target sequence in the chromosome (Burt 2003). Homology-based DNA repair mechanisms of the cell recognize the cut ends and fill in the gap by copying from the homologous chromosome. Thus in a heterozygote for the homing endonuclease gene, if the endonuclease is encoded opposite the cut site, the cell will repair the cut by copying the endonuclease gene into the other chromosome, in essence turning the heterozygote into a homozygote. Given this simple mechanism, it initially seemed that any suppression homing endonuclease gene would quickly select resistance in the form of a mutation in the target sequence (Burt 2003; Unckless *et al.* 2015, 2017; Drury *et al.* 2017). Resistance would then be allelic to the drive. However, this type of resistance now seems generally avoidable by designing CRISPR endonucleases to cut multiple sites within the target gene, only one of which needs to be wild-type for the drive to operate (Champer *et al.* 2018; Oberhofer *et al.* 2018). Other types of resistance to suppression drives are possible in principle, such as micro RNAs that block expression and function of CRISPR, inbreeding, and proteins that interfere with the CRISPR complex (Bull 2016; Stanley and Maxwell 2018). These various mechanisms will not typically be allelic to the drive and may even be fully unlinked (Lyttle 1981; Champer *et al.* 2019). Some may also entail a (large) fitness cost (Lyttle 1981).

As shown here, the evolution of non-allelic resistance can be avoided with drives that impose only a modest fitness cost on the organism, provided resistance is not initially common in the population. Drives that push population fitness to zero, which are tempting because of their assured population extinction (e.g., Lyttle 1977; Kyrou *et al.* 2018) will select resistance – but only if resistance arises (Lyttle 1979, 1981). Non-allelic resistance evolves by genetic hitchhiking from (negative) linkage disequilibrium with the drive locus. If the drive allele can evolve quickly to fixation then *D* will decay rapidly to zero, precluding further increases in resistance. This finding appears to be general, transcending 1-sex and 2-sex homing drives and toxin-antidote systems, with various assumptions about dominance and fitness effects. There are obviously many other combinations of effects to analyze, but a gene drive evading non-allelic resistance is clearly a possibility for many systems.

One ramification of these findings is that modification drives with unintended (mild) fitness consequences will not likely fail because of resistance evolution. Other types of evolution may still thwart modification drives, such as selection against their cargo. But formal resistance should not be a problem. However, what about suppression drives? Given that any single drive capable of assuredly evading resistance would provide only modest levels of population suppression, a single resistance-free drive might have only trivial effects on population numbers. Therefore, how feasible is meaningful population suppression by this approach?

The most assured design for a single partial-suppression drive is to target it to destroy a gene that is haplo-sufficent but compromises fitness moderately (not severely) in null homozygotes. Population extinction would likely require a succession of such gene drives, each targeting different genes, although simultaneous release multiple drives may offer a solution. (Whether simultaneous mild drives collectively evade resistance is an outstanding question.) Choose a target gene with too strong of a fitness effect, and the drive will not fix and may well select resistance – thwarting all future gene drives that work by the same mechanism. Choice of a target gene with appropriate fitness effects may be relatively easy for the first drive release but may increasingly require extensive background work as the organism’s fitness declines from previous drive fixations; and resistance levels may ratchet up with each release. Nonetheless, various strategies might be employed to enhance the long term effect of a single drive, such as choosing a target gene with a delayed or gradual effect on population fitness. Genes important in defense against an initially uncommon predator or important in surviving an environmental factor that can be controlled may be possibilities, as does destroying genes to reduce economic impact without impairing other aspects of the life history. The genetic effect of a drive is sudden, but the ecological effect can be gradual or specific.

### Spillover and containment

The findings here bear on concerns about the potential of gene drives to escape from target populations and do unintended harm. Any single suppression drive with mild fitness consequences may be introduced to reduce the economic or public health burden of a pest without causing pest extinction. Escape of a ‘mild’ drive into other species is also expected to be less consequential than is escape of an extremely deleterious drive. And drives that have only modest fitness effects may feasibly target genes that are poorly conserved between species, reducing their likelihood of escape. However, whatever genomic target is chosen, it must be sufficiently invariant that the drive can fix throughout the target populations of a species, lest allelic resistance evolve.

A downside of the findings here is that engineered resistance to a gene drive (such as resistance using known anti-CRISPR proteins) need not result in drive suppression. Engineered resistance seems like an obvious fail-safe against unwanted escape, and it no doubt is an assured defense against highly deleterious drives. But drives with modest fitness effects may not be sufficient to select engineered resistance unless the resistance can be introduced at high levels or the resistance locus can be tightly linked to the drive.

Alternative gene drive designs are being considered to reduce unintended spread. Some of those designs require introduction above a threshold frequency, such that low levels of migration would not be sufficient for them to spread beyond the release population (Vella *et al.* 2017; Dhole *et al.* 2018). It remains to be shown whether drives that are easily contained will be susceptible to resistance evolution; some containment strategies apparently rely on large fitness effects to minimize risk of escape (Tanaka *et al.* 2017; Greenbaum *et al.* 2019), which could make them tend to select resistance.

### Caveats

Fixation of a gene drive does not necessarily end the evolutionary response to a gene drive. Annihilation of gene function throughout a species by a gene drive may select compensatory evolution in other genes, ameliorating the impact (Burt 2003). Furthermore, any off-target activity of CRISPR may select suppression of CRISPR activity (Jeong *et al.* 2020) even though the drive has fixed, thereby limiting opportunities to introduce additional drives. Implementations of resistance-free drives will thus need to be monitored for long term effects.

As with most studies of gene drive resistance, our models assumed fully mixed populations, a state especially conducive to gene drive spread. Population structure, a key component of many natural populations, will retard gene drive spread and may even qualitatively alter outcomes. It will be interesting to extend the present analyses to population structure to see how selection of resistance is altered (North *et al.* 2013; *Beaghton et al.* 2016; North *et al.* 2019; Champer *et al.* 2020b). Minimally, we expect structure to alter the dynamics of linkage disequilibrium that is key to the evolution of resistance. Inbreeding, as one form of structure, is known to hamper the spread of suppression drives (Bull 2016; Bull *et al.* 2019) directly but also impacts the process of genetic hitchhiking (Hedrick 1980) which could indirectly favor the rise of non-allelic resistance. It may also affect the evolution of other types of resistance.

## Acknowledgments

We thank Bruce Hay and Jackson Champer for comments. Comments by two reviewers and L. Wahl on the first submitted draft led us to greatly expand the analytical and numerical analyses. RG was supported by a WSU Honors College Distinguished Professorship, MLT by a WSU College of Arts and Sciences Undergraduate Research grant, and JJB by NIH grant GM 122079.

## Appendix Derivations

## Effects of transmission

In this section, we show how transmission affects the population genetic structure of the drive and resistance loci. Consider a diploid population in which the frequency of genotypes with *i* copies of *A* and *j* copies of *R* at the start of meiosis is *Z*_*ij*_ for *i*, *j* = 0, 1, 2 and the frequency of haplotypes with *k* drive allele *A* and *l* resistance alleles *R* is *x*_*kl*_ for *k*, *l* = 0, 1. The allele frequencies and linkage disequlibrium are, respectively, *p*_*A*_ = 1 − *p*_*a*_ = *x*_10_ + *x*_11_, *p*_*R*_ = 1 − *p*_*r*_ = *x*_01_ + *x*_11_, and *D* = *x*_00_*x*_11_ − *x*_01_*x*_10_. Note that we use a different notation for the haplotype frequencies in the main text that is easier to grasp but less convenient for derivations; the equivalencies are as follows: *x*_00_ ↔ *x*_*ar*_, *x*_01_ ↔ *x*_*aR*_, *x*_10_ ↔ *x*_*Ar*_, and *x*_11_ ↔ *x*_*AR*_.

We assume the gene drive and resistance loci are linked with recombination rate 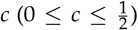 and that the amount of segregation distortion depends on the resistance locus genotype. A genotype with resistance genotype *G* (*G* = *rr*, *Rr*, *RR*) distorts segregation such that a drive locus heterozygote produces 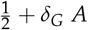 and 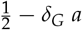 alleles, where *δ*_*G*_ is the distortion parameter 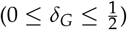. Complete distortion occurs if 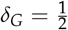; *δ*_*G*_ = 0 indicates no distortion (i.e., equal segregation).

After transmission, the frequencies of gametes produced by the diploids are

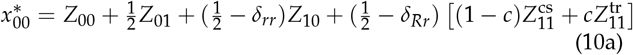

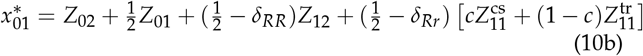

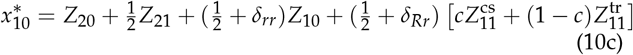

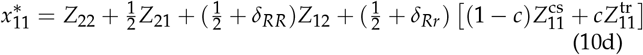

where 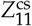 and 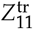 are the frequencies of ‘cis’ (*AR*/*ar*) and ‘trans’ (*Ar*/*aR*) conformation double heterozygotes, respectively. Using these equations, it can be shown that

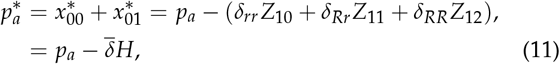

where *H* = *Z*_10_ + *Z*_11_ + *Z*_12_ and 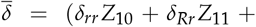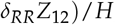, and

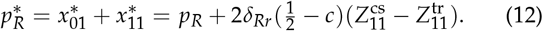

If we now assume the population was formed by random union of gametes, then

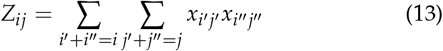

with 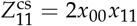 and 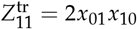. In this case,

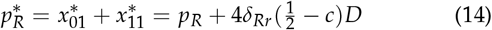

and the gamete recursions are equivalent to

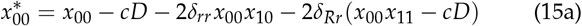

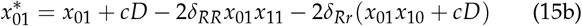

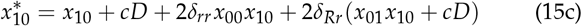

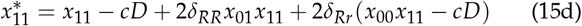

Equations (15) can also be used to show that disequilibrium after transmission is

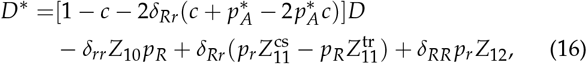

where

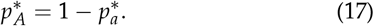

## Model of costly resistance: gametic selection

Consider the model described in the previous subsection but assume multiplicative fitnesses at both the drive and resistance loci such that each copy of the drive allele *A* reduces an individual’s fitness by the multiplicative factor 1 − *s*_*A*_ and each copy of the resistance allele *R* reduces fitness by the factor 1 − *s*_*R*_. This is called gametic selection (e.g., Lewontin 1970). The evolutionary dynamics of independent loci with gametic selection are mathematically equivalent those resulting from diploid multiplicative fitnesses (e.g., Crow and Kimura 1970).

Using (14), (15), (16), and (17), selection after transmission results in the following haplotype frequencies at the start of the next generation:

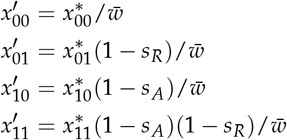

where

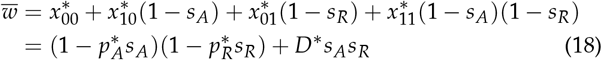

These expressions can be used to show that the corresponding frequencies of the drive and resistance alleles are

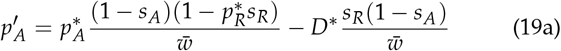

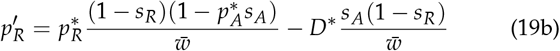

If resistance has no cost (*s*_*R*_ = 0) and is complete and dominant (i.e., *δ*_*RR*_ = *δ*_*Rr*_ = 0), equations (19) are equivalent to equations (2) and (6). However, selection against resistance, *s*_*R*_ > 0, could indirectly promote the drive allele *A* if the loci have a negative association (second term in eq. 19a). For 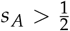, it is possible to prove that this model has an unstable equilibrium with no resistance 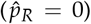 and polymorphism for the drive

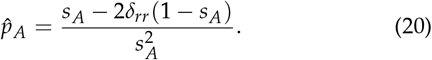

## Model of a lethal gene drive with linked resistance

In this subsection, we describe a model similar to the above except that the gene drive is recessive lethal. Assuming death occurs after diploids are formed by random mating as in (13) but before transmission, the post-death frequencies (indicated by the symbol #) are

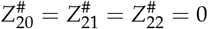

and, for *i* = 0, 1,

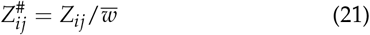

where

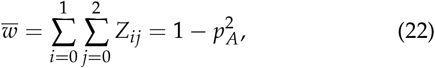

the fraction surviving. From this it can be shown that

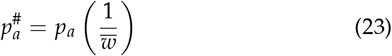

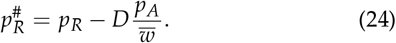

Equation (24) shows that when *D* < 0 the benefit to resistance from lethality increases with the frequency of the drive allele.

Applying the transmission equations (10) to the post-death zygote frequencies 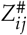 and assuming resistance is complete and dominant (*δ*_*RR*_ = *δ*_*Rr*_ = 0), transmission from the surviving genotypes produces the following gamete frequencies:

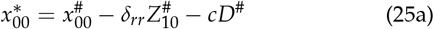

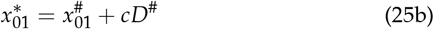

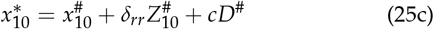

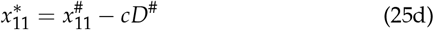

where gamete frequencies among the survivors are

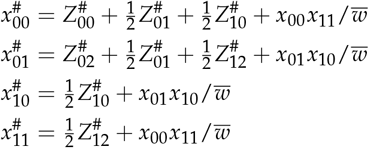

and

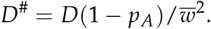

Note from (25) that

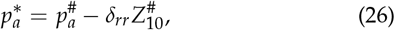

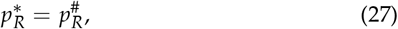

and

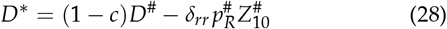

where 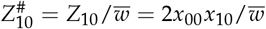 is the frequency of resistance-free heterozygotes among survivors. Equations (27) and (24) show that even with a recessive lethal gene drive, the evolutionary spread of perfect, dominant resistance is completely determined by genetic hitchhiking with the wild-type drive allele during selection.

